# Ubiquitous lognormal distribution of neuron densities in mammalian cerebral cortex

**DOI:** 10.1101/2022.03.17.480842

**Authors:** Aitor Morales-Gregorio, Alexander van Meegen, Sacha J. van Albada

**Affiliations:** Institute of Neuroscience and Medicine (INM-6) and Institute for Advanced Simulation (IAS-6) and JARA-Institut Brain Structure-Function Relationships (INM-10), Jülich Research Centre, Jülich, Germany; Institute of Zoology, University of Cologne, Cologne, Germany

**Keywords:** neuron density, cytoarchitecture, lognormal distribution, neurogenesis, mathematical model

## Abstract

Numbers of neurons and their spatial variation are fundamental organizational features of the brain. Despite the large corpus of cytoarchitectonic data available in the literature, the statistical distributions of neuron densities within and across brain areas remain largely uncharacterized. Here, we show that neuron densities are compatible with a lognormal distribution across cortical areas in several mammalian species, and find that this also holds true within cortical areas. A minimal model of noisy cell division, in combination with distributed proliferation times, can account for the coexistence of lognormal distributions within and across cortical areas. Our findings uncover a new organizational principle of cortical cytoarchitecture: the ubiquitous lognormal distribution of neuron densities, which adds to a long list of lognormal variables in the brain.

## Introduction

Neurons are not uniformly distributed across the cerebral cortex; their density varies strongly across areas and layers (Brodmann, 1909, von Economo et al., 2008). The neuron density directly affects short-range as well as long-range neuronal connectivity (Braitenberg and Schüz, 1991, Ercsey-Ravasz et al., 2013). Elucidating the distribution of neuron densities across the brain therefore provides insight into its connectivity structure and, ultimately, cognitive function. Additionally, statistical distributions are essential for the construction of computational network models, which rely on predictive relationships and organizational principles where the experimental data are missing (Hilgetag et al., 2019, van Albada et al., 2022). Previous quantitvative studies have provided reliable estimates for cell densities across the cerebral cortex of rodents (Charvet et al., 2015, Erö et al., 2018, Herculano-Houzel et al., 2013), non-human primates (Atapour et al., 2019, Beul and Hilgetag, 2019, Charvet et al., 2015, Collins et al., 2010, 2016, Turner et al., 2016), large carnivores (Jardim-Messeder et al., 2017), and humans (von Bartheld et al., 2016, von Economo et al., 2008). However, to the best of our knowledge, the univariate distribution of neuron densities across and within cortical areas has not yet been statistically characterized. Instead, most studies focus on qualitative and quantitative comparisons across species, areas, or cortical layers. Capturing the entire distribution is necessary because long-tailed, highly skewed distributions are prevalent in the brain (Buzsáki and Mizuseki, 2014) and invalidate the intuition—guided by the central limit theorem—that the vast majority of values are in a small region of a few standard deviations around the mean.

Here, we characterize the distribution of neuron densities *ρ* across mammalian cerebral cortex. Based on the sample histograms (Figure 1) we hypothesize that *ρ* follows a lognormal distribution, similar to many other neuroanatomical and physiological variables (Buzsáki and Mizuseki, 2014) such as synaptic strengths (Robinson et al., 2021), synapse sizes (Dorkenwald et al., 2022, Loewenstein et al., 2011, Santuy et al., 2018), axonal widths (Liewald et al., 2014, Wang et al., 2008), and cortico-cortical connection densities (Gămănuţ et al., 2018, Markov et al., 2014). We used neuron density data from mouse (*Mus musculus*), marmoset (*Callithrix jacchus*), macaque (*Macaca mulatta*), human (*Homo sapiens*), galago (*Otolemur garnettii*), owl monkey (*Aotus nancymaae*), and baboon (*Papio cynocephalus anubis*) to test this hypothesis (see Cell density data for a detailed description of the data). The marmoset, galago, owl monkey, baboon, and macaque_2_ data sets are based on a single subject; the mouse, macaque_1_, and human data sets are based on a combination of data across several subjects. The statistical tests conclude that the hypothesis cannot be rejected in the majority of cases, suggesting that the underlying distribution is compatible with a lognormal distribution if the samples are based either on cytoarchitectonically defined areas or on uniformly sampled regions. Beyond the distribution across cortical areas, we show that neuron densities within most areas of marmoset cortex are also compatible with a lognormal distribution. To complement the statistical tests, we perform a model comparison with several other distributions and find that none outperform the lognormal distribution as a model of the data within and across areas. Finally, we show that the lognormal distribution within cortical areas can emerge during neurogenesis from a simple cell division model with variability. The model can furthermore account quantitatively for the lognormal density distribution across areas based on an inferred distribution of proliferation times of the areas. Additional between-area variability in the proliferation rates is also compatible with the model but not necessary to obtain a quantitative agreement with the observed distribution. Thus, our model shows how the lognormal distribution of neurons could emerge both within and across cortical areas.

**Figure 1:**
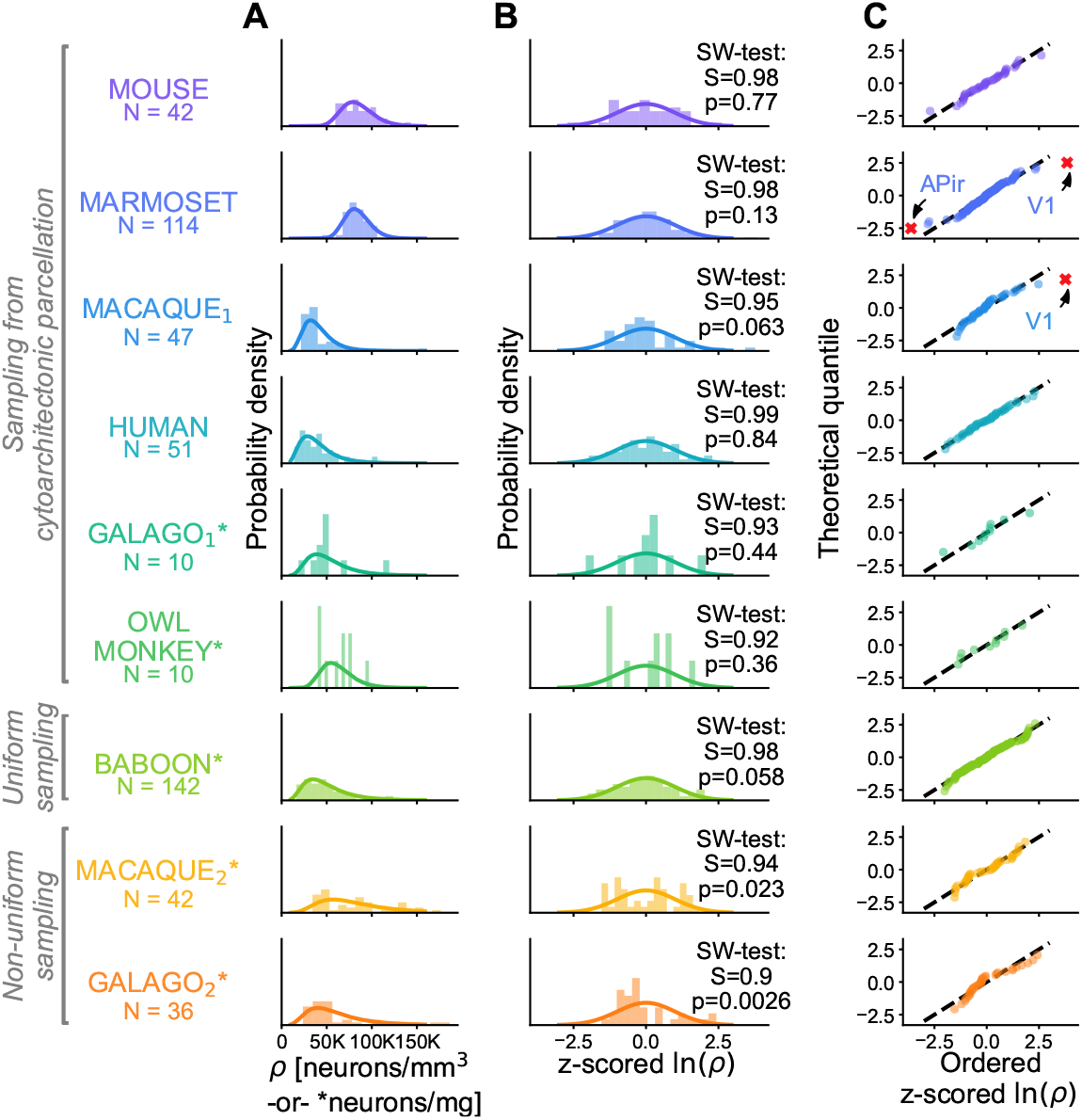
Neuron and cell densities *ρ* follow a lognormal distribution across cortical areas for multiple species. Different data sets for the same species are denoted with subscript indices (see Cell density data). **A** Histogram of *ρ* (bars) and probability density function of a fitted lognormal distribution (line). The number of samples *N* is either the number of sampled cytoarchitectonic areas (mouse, marmoset, macaque_1_, human, galago_1_, and owl monkey) or the number of sampling frames (baboon, macaque_1_, and galago_2_). For marmoset, galago, owl monkey, baboon, and macaque_2_ the data is based on a single subject; for mouse, macaque_1_, and human it is based on a combination of data across several subjects. **B** Z-scored ln(*ρ*) histogram (bars), standard normal distribution (line), and result of the Shapiro-Wilk normality test. **C** Probability plot of z-scored ln(*ρ*). Discarded outliers marked with a red cross.

## Materials and methods

### Cell density data

Estimates of neuron density for the available cortical areas across the mouse (*Mus musculus*), marmoset (*Callithrix jacchus*), macaque (*Macaca mulatta*), human (*Homo sapiens*), galago (*Otolemur garnettii*), owl monkey (*Aotus nancymaae*), and baboon (*Papio cynocephalus anubis*) cerebral cortex were used in this study.

In the cases of mouse, marmoset, macaque_1_, human, galago_1_, and owl monkey the data were sampled from standard cytoarchitectonic parcellations; abbreviated names for all areas are listed in Table S1. Note that we use subscript indices to distinguish between different data sets on the same model animal, e.g., macaque_1_ and macaque_2_.

Neuron density estimates for the mouse were published by Erö et al. (2018), and were measured from previously published Nissl-body-stained slices (Lein et al., 2007), where genetic markers were used to distinguish between cell types. The data were provided in the Allen Brain Atlas parcellation (Dong, 2008, Lein et al., 2007). To estimate the quantitative neuron and cell densities Erö et al. (2018) used “a variety of whole brain image datasets [from the Allen Mouse Brain Atlas]” as well as “some values reported from anatomical experiments in the literature”, such as from (Herculano-Houzel et al., 2013).

Neuron density estimates for the marmoset cortex were published by Atapour et al. (2019), and were measured from NeuN-stained slices. The data were provided in the Paxinos parcellation (Paxinos et al., 2011) and all quantitative values are derived from the same brain of a single subject. Neuron densities within each counting frame used in the original publication (Atapour et al., 2019, their Figure S1) were obtained via personal communication with Nafiseh Atapour, Piotr Majka, and Marcello G. Rosa.

The neuron density estimates in the first macaque data set, macaque_1_, were previously published in visual form by Beul and Hilgetag (2019), and were obtained from both Nissl-body- and NeuN-stained brain slices. Most of the numerical values were reported by Dombrowski et al. (2001) and Hilgetag et al. (2016), and the remaining values were provided by Sarah F. Beul and Claus C. Hilgetag via personal communication. Counts based on Nissl-body staining were scaled according to a linear relationship with the counts from NeuN staining obtained from selected areas where both types of data were available (Beul and Hilgetag, 2019). The data follow the M132 parcellation (Markov et al., 2014) and constitute the average across subjects.

Cell density estimates for the human cortex were previously published by von Economo et al. (2008), and were measured from Nissl-body-stained brain slices. The human data therefore most likely reflect combined neuron and glia densities. The data were provided in the von Economo parcellation (von Economo et al., 2008); all quantitative values are based on “the mean of the numbers gathered from the various brains” (von Economo et al., 2008, p. 201).

Cell and neuron density estimates for galago_1&2_, owl monkey, baboon, and macaque_2_ were previously published by Collins et al. (2010), and were measured using the isotropic fractionator method. The data are sampled from common parcellation schemes in galago_1_ and owl monkey, approximately equal-size samples in the baboon, and irregular non-uniform samples in macaque_2_ and galago_2_. All the quantitative values were derived from one hemisphere of each subject separately. Each data set thus describes only a single subject, without combining data across individuals of the same species.

### Lognormality testing

To test for lognormality, we take the natural logarithm, ln(*ρ*), which converts lognormally distributed samples into normally distributed samples. We then test for normality of ln(*ρ*) using the Shapiro-Wilk (SW) test, the most powerful among a number of commonly applied normality tests (Razali and Yap, 2011). Large outliers (| z-scored ln(*ρ*) | ≥ 3) were excluded from the normality test; the excluded outliers are indeed cytoarchitectonically distinct areas (discussed below).

Note that any hypothesis test, including the SW test, cannot show that the distribution is lognormal. If the p-value is larger than a certain threshold one cannot reject the null hypothesis that the distribution of *ρ* is lognormal, i.e., the data is compatible with a lognormal distribution. Thus, we perform further tests, such as statistical model comparison, and comparing ln(*ρ*) across different animals with a Kolmogorov-Smirnov test.

### Statistical model comparison

In order to assess which model is most compatible with the data, we compared the relative likelihood of different distributions against each other. We included an extensive list of distributions with support on ℝ^+^, estimated the distributions’ parameters using maximum likelihood, and calculated the Akaike Information Criterion (*AIC*)

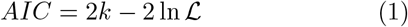

where *k* is the number of estimated parameters of the model and ℒ is the estimated maximum likelihood. We further compare the models using the relative likelihood (ℒ_*r*_)

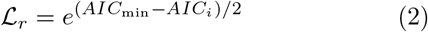

where *AIC*_min_ is the minimum *AIC* across all models and *AIC*_*i*_ is the *AIC* for the *i*th model. The relative likelihood indicates the probability that, from among the tested models, the *i*th model most strongly limits the information loss (Burnham and Anderson, 2004). We take a significance threshold of *α* = 0.05 on the relative likelihood to determine whether a model is significantly worse than the best possible model.

### Model of neurogenesis with variability

#### Within areas

Neurons are generated through symmetric or asymmetric cell division of neural progenitor cells (Cadwell et al., 2019). Grouping all types of progenitor cells into a single population *P*, all neurons into a population *N*, and excluding other post-mitotic cell types, their population size can be modeled by the coupled system of differential equations (Picco et al., 2018)

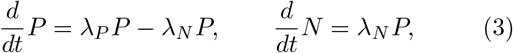

where *λ*_*P*_ denotes the (potentially time-dependent) rate of progenitor generation and *λ*_*N*_ the (potentially time-dependent) rate of neuron generation.

Following the radial unit hypothesis (Rakic, 2009), we consider a small number of such radial units (small compared to the total size of the area) and determine the density of progenitors *ρ*_*P*_ and neurons *ρ*_*N*_ in these radial units by dividing through the volume *V* of the considered radial units in the fully developed cortex. Importantly, this reference volume is the same for every area. Since Equation (3) is linear, dividing by *V* leads to

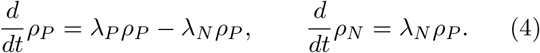

Note that this does not necessarily exclude tangential migration as long as the net influence of the tangential migration is zero.

#### Progenitor cell proliferation

First, we consider the proliferation of the progenitor cells, which we assume to be governed by a noisy rate. Modeling the noise by a zero-mean, unit-strength Gaussian white noise process *ξ*, we obtain a stochastic differential equation (SDE) for the progenitor cell density

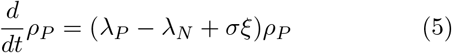

where *σ* controls the (potentially time-dependent) intensity of the noise. Using the Stratonovich interpretation— assuming that the noise process has a small but finite correlation time before taking the white-noise limit (Van Kampen, 2007)—the stochastic differential equation transforms by the same rules as an ordinary differential equation (Van Kampen, 2007) such that we can rewrite the SDE (5) as 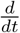 ln *ρ*_*P*_ = *λ*_*P*_ − *λ*_*N*_ + *σξ* with the solution

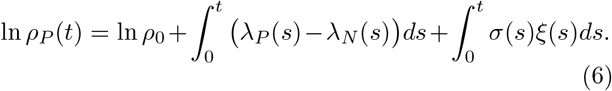

Since *ξ*(*t*) is Gaussian and Equation (6) is linear, ln *ρ*_*P*_ (*t*) is Gaussian and hence *ρ*_*P*_ (*t*) is lognormally distributed at all times *t*. The parameters of this lognormal distribution are the mean of the logarithmic progenitor cell density *μ*_*P*_ (*t*) = ⟨ln *ρ*_*P*_ (*t*)⟩ and the variance of the logarithmic progenitor cell density *σ*_*P*_ (*t*)^2^ = ⟨Δ(ln *ρ*_*P*_ (*t*))^2^⟩ (here Δ*x* ≡ *x* − ⟨ *x* ⟩). Using Equation (6), ⟨ *ξ*(*s*) ⟩ = 0, and ⟨ *ξ*(*s*)*ξ*(*s*^′^)⟩ = *δ*(*s* − *s*^′^), we obtain (cf., for instance Braumann (2007))

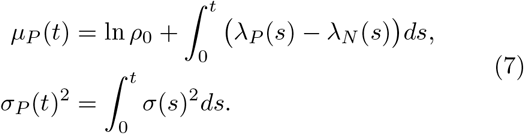

Thus, the progenitor cell densities are lognormally distributed at all times with parameters *μ*_*P*_ (*t*) and *σ*_*P*_ (*t*). The corresponding first two moments of the progenitor cell density are

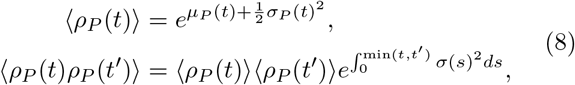

where we used the characteristic functional of the Gaussian white noise *ξ*(*t*) (Van Kampen, 2007),

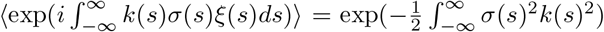, with the test function *ik*(*s*) = 𝟙 _[0,*t*]_(*s*) for the first moment (here 𝟙 _*A*_(*s*) denotes the indicator function) and test function *ik*(*s*) = 𝟙 _[0,*t*]_(*s*) + 𝟙 _[0,*t*]_(*s*) for the sec-ond moment as well as 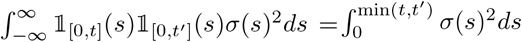.

#### Neurogenesis

Next, we consider neurogenesis. We assume that the noise affects primarily the rate of progenitor cell proliferation. The solution to Equation (4) for a given *ρ*_*P*_ (*t*) and initial condition *ρ*_*N*_ (0) = 0 is

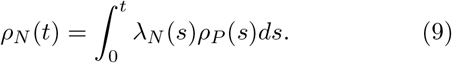

Since *ρ*_*N*_ (*t*) is the integral of the (temporally correlated) lognormal process *ρ*_*P*_ (*s*) it is formally not lognormal. However, the sum of independent lognormal random variables is well approximated by a lognormal distribution with matched first and second moment (Fenton, 1960, Marlow, 1967). Here, we extend this approximation to the integral of temporally correlated lognormal processes. The first two moments of the neuron density follow from the averages of Equation (9),

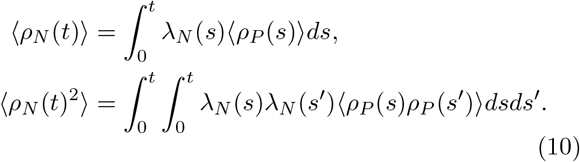

The lognormal approximation with matched moments is parameterized by

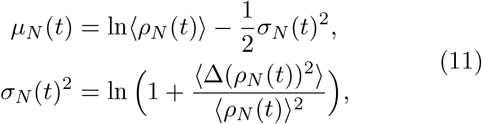

where we used 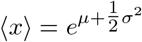 and 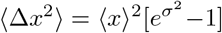 for a lognormal variable *x*. Note that the parameters of the lognormal distribution are the mean of the logarithmic neuron density *μ*_*N*_ (*t*) = ⟨ln *ρ*_*N*_ (*t*)⟩ and the variance of the logarithmic neuron density *σ*_*N*_ (*t*)^2^ = ⟨Δ(ln *ρ*_*N*_ (*t*))^2^⟩.

#### Across areas

Thus far, the model accounts for the lognormal distribution of neuron densities within an area. Across areas, we hypothesize that the distribution of proliferation times (Cadwell et al., 2019, Rakic, 2002) is the most important cause of the variability in the neuron densities. To characterize an individual area, we consider the average density ⟨*ρ*_*N*_ (*t*) ⟩. For convenience, we introduce the auxiliary quantity

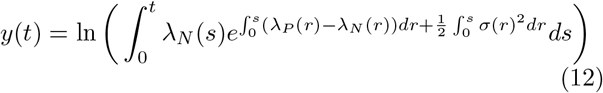

such that the logarithm of the mean neuron density simplifies to

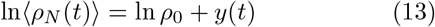

where we inserted Equation (7) into Equation (8) to obtain the mean of the progenitor cell density, which we then plugged into Equation (10).

In order to arrive at a lognormal distribution of the mean density ⟨ *ρ*_*N*_ (*t*) ⟩ across areas, the terms on the r.h.s. of Equation (13) have to be normally distributed. In particular, this means that the proliferation times have to be distributed such that *y*(*t*) is Gaussian. The distribution *p*(*y*) of *y* = *y*(*t*), which is a strictly monotonic transformation of the proliferation time *t*, is related to the distribution of proliferation times *p*(*t*) through (Van Kampen, 2007)

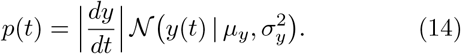

The first factor on the r.h.s. can be obtained from Equation (12) and the second factor is a Gaussian probability density with mean *μ*_*y*_ and variance 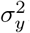. Hence, Equation (14) fully specifies the distribution of proliferation times; conversely, for proliferation times distributed according to Equation (14), the mean neuron density ⟨*ρ*_*N*_ (*t*) ⟩ is lognormally distributed across areas. Note that since the Gaussian has support on the entire real line, the neuron density needs to diverge for *t* → ∞. Thus, we restrict *λ*_*P*_ (*t*), *λ*_*N*_ (*t*), and *σ*(*t*) such that the neuron density would diverge in the hypothetical limit of an infinite proliferation time.

#### Parameter estimation

Above, we showed how a noisy rate of progenitor proliferation leads to a lognormal distribution of progenitor cell densities and neuron densities within an area and, for the distribution of proliferation times (14), to a lognormal distribution of (within-area) mean neuron densities across areas. In order to compare the predictions of the model with the data, we estimate the model’s parameters using the available experimental data.

We restrict the analysis to the simplified case of constant rate *λ*_*P*_, *λ*_*N*_, and noise intensity *σ* and identical rates of progenitor and neuron proliferation, *λ*_*P*_ = *λ*_*N*_ ≡ *λ*. In particular the assumption of a constant rate (note that this is not a necessary assumption for the above theory) is simplifying because the cell cycle length varies during development (Kornack and Rakic, 1998). Despite this simplifying assumption, the model quantitatively matches the data for the parameters inferred below (Figure 4).

First, we determine the rate *λ* from the average cell cycle length of progenitor cells determined from a short period of 2 hours (Kornack and Rakic, 1998) such that fluctuations and the conversion of progenitor cells to neurons can be neglected. For a given cell cycle length ℓ, the number of cells increases as 2^*t/*ℓ^ = exp *t* ln(2)*/*ℓ. Thus, the cell cycle length ℓ corresponds to a proliferation rate *λ* = ln(2)*/*ℓ. Using the average cell cycle length of ℓ ≈ 1.5 days from macaque (Kornack and Rakic, 1998, Picco et al., 2018), we obtain *λ* ≈ 0.46 days^−1^. The proliferation time of areas varies between 30 and 60 days in macaque (Rakic, 2002) which we expect to be similar in the marmoset since macaques and marmosets have similar gestation times of 5.5 and 4.5 months, respectively (Schultz-Darken et al., 2016). Thus, we set the median proliferation time per area to *t*_1*/*2_ = 45 days, which determines *μ*_*y*_ = *y*(*t*_1*/*2_) because the median of the distribution (14) is given by 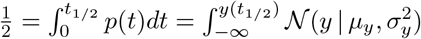 and the me-dian of the normal distribution is at *y* = *μ*_*y*_.

In addition, the mean *μ*_*y*_ of the auxiliary variable *y* is also constrained by the distribution of the variance 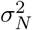 of the logarithmic neuron density across areas; we will use this additional constraint to determine the noise intensity *σ*. For a fixed proliferation time *t, σ*_*N*_ (*t*)^2^ is given by Equation (11). Since 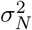 is a strictly monotonically increasing function of *t*, its distribution can be written in terms of that of the inverse 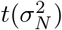, i.e., 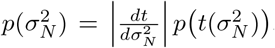. Using Equation (14) for the distribution of proliferation times *p*(*t*) and Equation (12) to relate *y*(*t*) and *t*, the distribution of 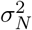 across areas is thus

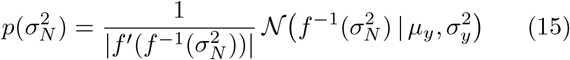

where 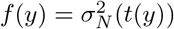 with *t*(*y*) the inverse of

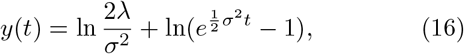

which follows from evaluating Equation (12) with *λ*_*P*_ = *λ*_*N*_ ≡ *λ* and constant *σ*^2^, and 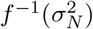 is the inverse of *f* (*y*). Explicitly, it is given by

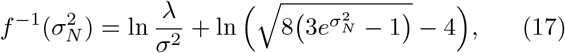

which follows from inserting *t*(*y*) into 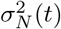 determined by Equation (11) and solving it for *y*. The maximum likelihood estimator 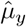 for *μ*_*y*_ with the likelihood 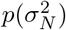 is the empirical average of 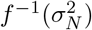 across areas because the Jacobian 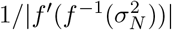 in Equation (15) does not depend on the parameters and hence does not affect the Gaussian likelihood of *μ*_*y*_. Using the marmoset data, we obtain 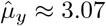. We finally arrive at *σ* ≈ 0.061 days^−1*/*2^ by enforcing 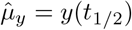, where *y*(*t*) is given by Equation (16), with *t*_1*/*2_ = 45 days. With *σ* and *μ*_*y*_ determined, we choose *σ*_*y*_ such that less than 2% of the proliferation times are smaller than 30 days or larger than 60 days. This leads to 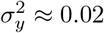.

It remains to estimate the initial progenitor density *ρ*_0_. To this end, we consider the distribution of ln⟨*ρ*_*N*_ ⟩. For a fixed proliferation time *t*, ln ⟨*ρ*_*N*_ (*t*) ⟩ is determined by Equation (13). Thus, it is also a monotonic transformation of the random variable *t*, with distribution 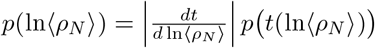. Using the linear dependence between ln ⟨ *ρ*_*N*_⟩ and *y*, Equation (13), in combination with the distribution of proliferation times (14), we obtain

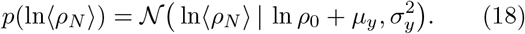

The maximum likelihood estimator for ln *ρ*_0_ + *μ*_*y*_ is the empirical average of ln ⟨*ρ*_*N*_⟩ across areas; subtracting *μ*_*y*_ we obtain *ρ*_0_ ≈ 3.8 × 10^3^ cells*/*mm^3^.

Note that in the simplified case considered here, the resulting distribution of proliferation times approaches a lognormal distribution in the left tail and a Gaussian in the right tail: for small times, 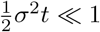, a Taylor expansion of *y*(*t*) leads to *y*(*t*) ≈ ln(*λt*) and thus a lognormal distribution; for large times, 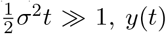 grows lin-early with 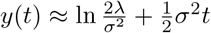 and thus the distribution approaches a Gaussian.

#### Simulation Details

We solve the SDE (5) using the Euler-Maruyama method with time step Δ*t* = 0.05 in total *N*_sample_ times with identical parameters for each of the 114 areas. Because the number of samples per area *N*_sample_ varies, we randomly choose *N*_sample_ for each area following *N*_sample_ ∼ Poisson(36.6) where 36.6 is the average sample size per cortical area in the marmoset data. The subsequent analysis of the model data is identical to the analysis of the experimental data.

## Results

### Lognormal distribution of neurons across cortical areas

We consider the neuron density distribution across cortex for several species (see Cell density data). The Shapiro-Wilk (SW) test (see Lognormality testing) concludes that the normality hypothesis of ln(*ρ*) cannot be rejected for mouse, marmoset, macaque_1_, human, galago_1_, owl monkey, and baboon (Figure 1B). For the data sets macaque_2_ and galago_2_ the normality hypothesis is rejected (*p* < 0.05); however, in these data sets, the densities were sampled neither uniformly nor based on a cytoarchitectonic parcellation. The normality hypothesis for the distribution of logarithmic densities across cytoarchitectonic areas is further supported by Figure 1C, which shows that the relation between theoretical quantiles and ordered samples is almost perfectly linear except for macaque_2_ and galago_2_.

For lognormality testing we removed the large outliers (marked with a red cross in Figure 1C). The outliers are area V1 in macaque_1_ and marmoset, which have densities far outside the range for all other areas in both species, and area APir in marmoset, which has a noticeably distinct cytoarchitecture with respect to the rest of the cerebral cortex (Atapour et al., 2019).

Next, we test the z-scored ln(*ρ*) from the different species and data sets against each other and find that they are pairwise statistically indistinguishable (*α* = 0.05 level; two-sample two-sided Kolmogorov-Smirnov test, see Figure S1 for full test results).

Additionally, we control for cell types in the distributions of the mouse, galago_1_, owl monkey, and baboon data. In the mouse data, different types of neurons and glia were labeled with specific genetic markers and their respective densities were reported separately for all cell types (Erö et al., 2018). In the galago_1_, owl monkey, and baboon data sets, the total numbers of cells and neurons were reported separately (Collins et al., 2010). We show that the neuron density distributions for all subtypes of neurons in the mouse are compatible with a lognormal distribution (Figure S2; SW test on ln(*ρ*), *p* > 0.05) while glia are not—with the notable exception of oligodendrocytes. When neurons and glia are pooled together (Figure S2C,D), the distribution of ln(*ρ*) still passes the SW normality test, likely due to the distribution being dominated by the neurons. Similar observations are made in the baboon data, where the glia do not pass the lognormality test, but the neurons do. In the cases of galago_1_ and owl monkey both the neurons and glia pass the lognormality test (Figure S2), which may, however, be partly due to the small number of density samples (N=12 in both cases). Thus, the mouse and baboon data—with large samples sizes (N=42 and N=142, respectively)—suggest that it is the neuron densities that follow a lognormal distribution but not necessarily the glia densities.

Finally, we perform a control test on the different staining types—Nissl and NeuN—using the macaque_1_ data. The staining methods differ in their treatment of glia: NeuN tends to label neuronal cell bodies only while Nissl indiscriminately labels both neurons and glia (Yurt et al., 2018). We show that regardless of staining type the cell densities pass the lognormality test (Figure S3; SW test on ln(*ρ*) with *p* > 0.05), suggesting that counting some glia in the cell densities does not confound our analysis of the macaque_1_ data.

Taken together, the normality test, the quantile plots, the pairwise tests, the cell type comparison, and the staining method comparison provide compelling evidence that the logarithmized neuron densities are normally distributed across cytoarchitectonic areas. This also holds for uniformly sampled neuron densities (baboon) but not for a sampling that is neither uniform nor based on a cytoarchitectonic parcellation (macaque_2_, galago_2_). Thus, the neuron densities are consistent with a lognormal distribution across the different cortical areas, as long as sampling is not irregular.

### Lognormal distribution of neurons within marmoset cortical areas

To investigate whether the lognormal distribution holds within cortical areas, we leverage detailed estimates of neuron density in marmoset (Atapour et al., 2019). Neurons were counted within 150 × 150 *μ*m counting frames for four strips per cortical area, all originating from the same subject. The within-area distributions of the sampled neuron densities *ρ*_*s*_ across the counting frames in three representative areas (MIP, V2, and V3; Figure 2A) again suggest a lognormal distribution. As before, we check for lognormality by testing ln(*ρ*_*s*_) for normality with the SW-test (for full test results see Table S2). At significance level *α* = 0.05, the normality hypothesis is not rejected for 89 out of 116 areas; whereas at *α* = 0.001, this is the case for 114 out of 116 areas (Figure 2B,C). Thus, regardless of the precise significance threshold, the lognormality hypothesis cannot be rejected within most cortical areas in the marmoset cortex.

**Figure 2:**
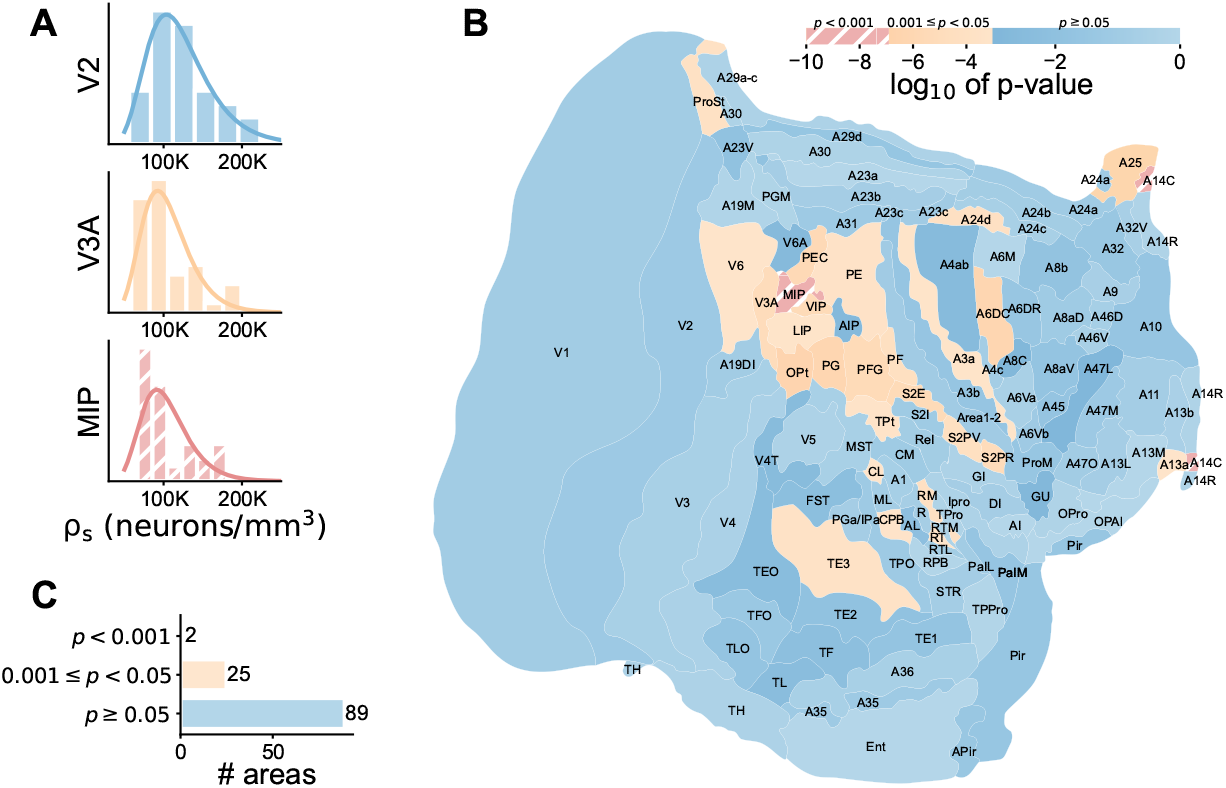
Neuron densities *ρ*_*s*_ follow a lognormal distribution within most areas of marmoset cortex. **A** Sample histograms of *ρ*_*s*_ and fitted lognormal distributions for three areas representing different degrees of lognormality. **B** Log_10_ of p-value of Shapiro-Wilk normality test of ln(*ρ*_*s*_) on a flattened representation of the marmoset cortex (Atapour et al., 2019). **C** Number of areas with p-values in the given significance ranges.

### Statistical model comparison

To complement the statistical hypothesis tests on the logarithmic densities, we compared the lognormal model with six other statistical distributions based on the relative likelihood (see Statistical model comparison). We included statistical distributions with support in R^+^ since neuron densities cannot be negative: lognormal, truncated normal, inverse normal, gamma, inverse gamma, Lévy, and Weibull. Of these, the lognormal, inverse normal, and inverse gamma distributions stand out as the distributions with the highest relative likelihoods, both across the entire cortex and within cortical areas (Figure 3A, Figure S4A). A visual inspection of the fitted distributions reveals that the lognormal, inverse nor-mal, and inverse gamma distributions produce virtually indistinguishable probability densities (Figure 3B, Figure S4C); indeed, the relative likelihoods of the three models are above 0.05 in all cases. This suggests that the data could theoretically be distributed according to either the lognormal, inverse normal, or inverse gamma distribution. To narrow down the model comparison, we show below how the lognormal distribution could arise from the simple biophysical process of noisy cell division. In contrast, we are not aware of a simple mechanism that could give rise to inverse normal or inverse gamma distributions.

**Figure 3:**
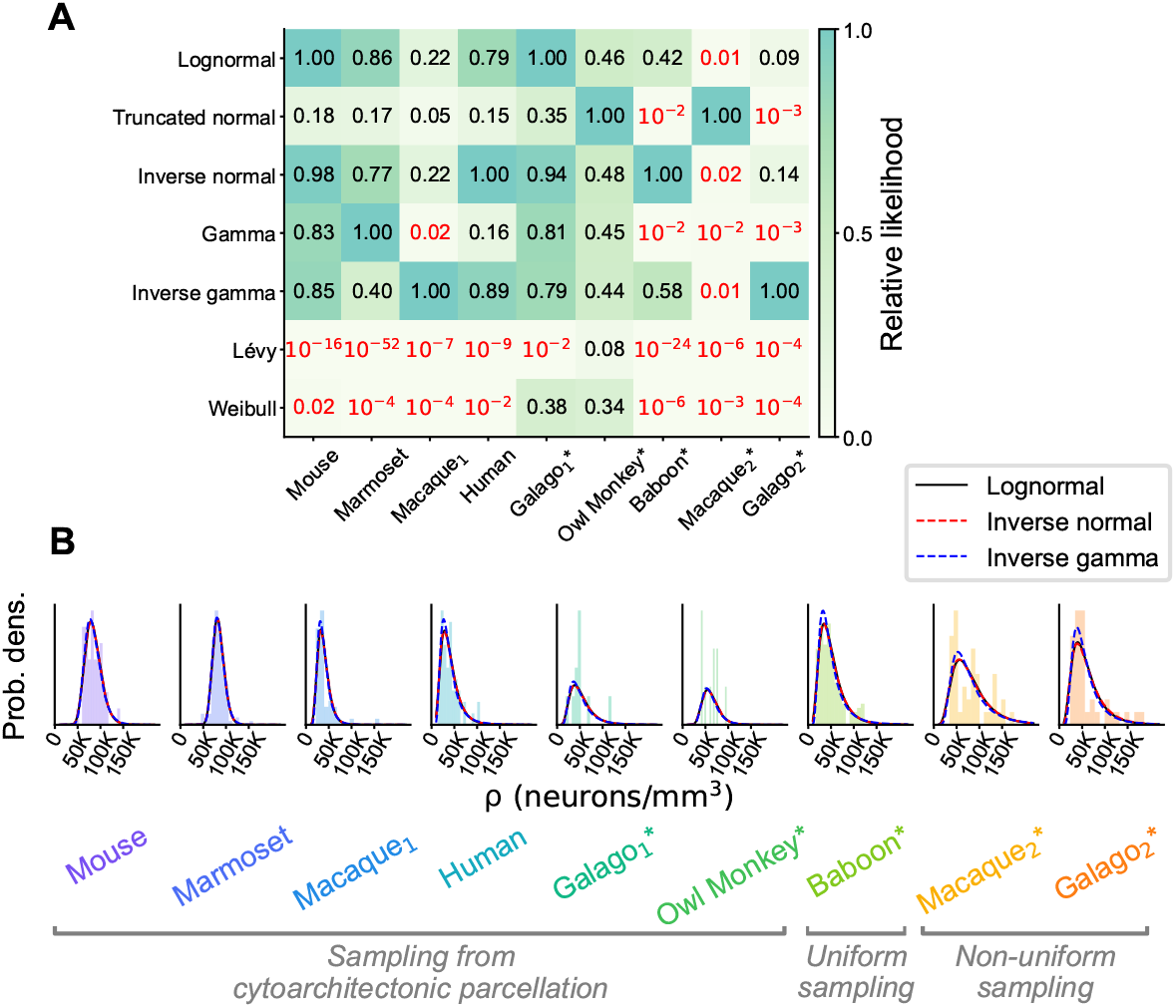
Statistical model comparison across the entire cortex of different animals. **A** Relative likelihood for seven compatible statistical models for all available area-level neuron density data sets; numerical values indicated for each model and animal. The red color indicates a relative likelihood < 0.05 with respect to the model with the highest likelihood. **B** The three best statistical models (according to the relative likelihood) fitted to the neuron density histograms in each animal; the three models produce visually nearly indistinguishable fits.

### A minimal model for the emergence of lognormally distributed neuron densities

The finding of lognormally distributed neuron densities raises the question how the intricate process of neurogenesis (Cadwell et al., 2019, Dehay and Kennedy, 2007, Rakic, 2009) culminates in this distribution within and across areas in several mammalian species.

On the within-area level, a minimal model shows that there is no need for a specific regulatory mechanism (see Model of neurogenesis with variability for further details): assuming that the proliferation of the neural progenitor cells is governed by a noisy rate *λ*_*P*_ (*t*) + *ξ*(*t*), where *λ*_*P*_ (*t*) denotes the mean rate and *ξ*(*t*) is a zero-mean Gaussian white noise, the resulting density of progenitor cells is lognormally distributed.

The cells produced by the cell division of the neural progenitor cells are terminally differentiated neurons, which thus do not divide further. This renders the final neuron density a linear integral of the (changing) neural progenitor cell density. While this additive process formally does not preserve the lognormality of the neural progenitor density, the resulting neuron density distribution is statistically indistinguishable from a lognormal distribution with matched moments (SW test *p* = 0.4 with *N* = 2000 samples in Figure S5F)—akin to the lognormal approximation for the sum of independent lognormal random variables (Fenton, 1960, Marlow, 1967). Put differently, the lognormally distributed density of neural progenitor cells leads to an approximately lognormally distributed density of neurons. Thus, the lognormal neuron density distribution within areas could be a hallmark of a progenitor cell proliferation process with variability.

On the across-area level, the proliferation times for each area become relevant, since they vary up to twofold (Rakic, 2002). We therefore hypothesize that the distribution of proliferation times is the most important source of the variability across areas (variability due to area-specific proliferation rates (Lukaszewicz et al., 2005) is discussed below). Since the average neuron density within an area is determined by the proliferation time, the distribution of proliferation times specifies the distribution of area-averaged neuron densities across areas. This relation can be inverted to determine the distribution of proliferation times from the lognormal distribution of (within-area) average neuron densities (see Model of neurogenesis with variability). A specific prediction of this model is that the logarithmic mean ln(⟨*ρ*⟩) and variance of ln(*ρ*) are related through the proliferation time—both mean and variability increase with proliferation time in approximate proportion to each other (see Equations (11) and (13), respectively). Indeed, we observe a linear correlation in the marmoset data (Pearson *r* = 0.51, *p* ≤ 10^−8^, Figure 4C) as well as in the data produced by the model (Pearson *r* = 0.4, *p* ≤ 10^−4^, Figure 4D).

**Figure 4:**
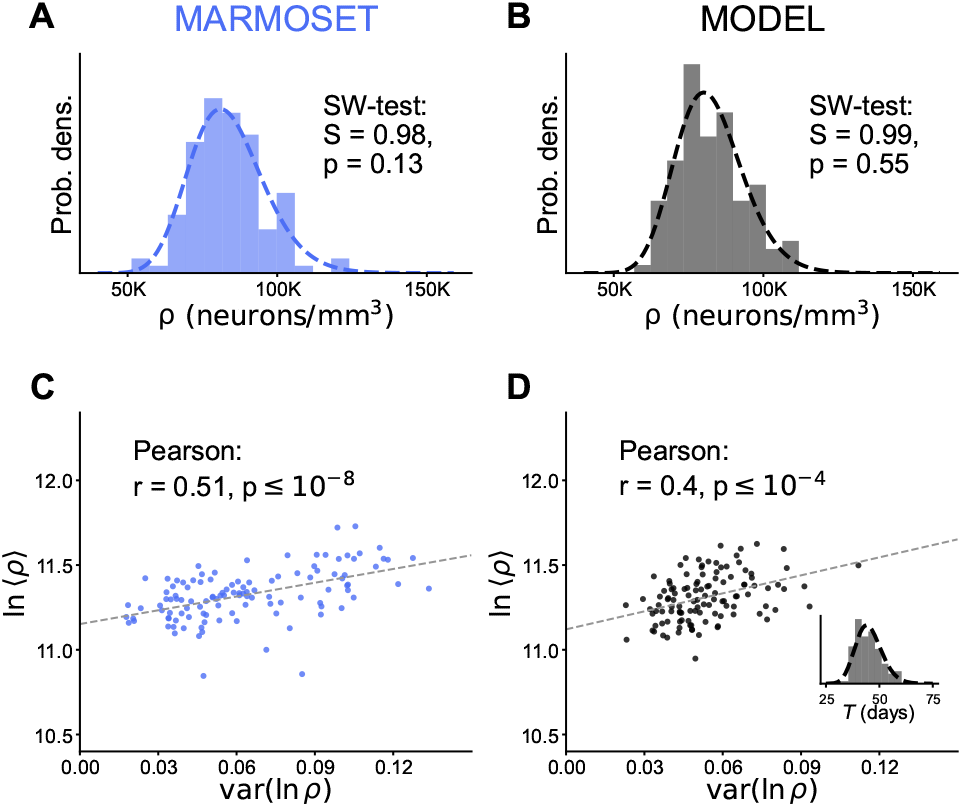
Neuron densities in the marmoset are compatible with a minimal model of neurogenesis with variability. **A, B** Distribution of neuron densities across areas for the marmoset (**A**) and the model (**B**). The Shapiro-Wilk test does not reject normality of ln(*ρ*) for both distributions. **C, D** The logarithmic mean ln(⟨*ρ*⟩) and the variance of the logarithmic density var(ln(*ρ*)) for each area are significantly correlated with each other (Pearson correlation test) for the marmoset (**C**) and the model (**D**). The inset in **D** displays the inferred distribution (dashed line; see Equation (14)) of proliferation times alongside the sample used for the simulation (bars). For all parameters of the model and further details see Model of neurogenesis with variability.

## Discussion

In this work we show that neuron densities are compatible with a lognormal distribution across cortical areas in multiple mammalian cortices and within most cortical areas of the marmoset, uncovering a ubiquitous organizational principle of cerebral cortex. The distributions of neuron and cell densities in general depend on the underlying spatial sampling: We found that neuron densities follow a lognormal distribution within cytoarchitectonically defined areas, across such areas, and when averaged within small parcels uniformly sampled across cortex, but not when sampled in a highly non-uniform manner not following cytoarchitectonic boundaries.

Furthermore, we show that none of a sizeable list of statistical models outperform the lognormal distribution. Our results are in agreement with the observation that surprisingly many characteristics of the brain follow lognormal distributions (Buzsáki and Mizuseki, 2014). Moreover, this analysis highlights the importance of characterizing the statistical distributions of brain data because simple summary statistics—such as the mean or standard deviation—lack nuance and are not necessarily a good representation of the underlying data.

These findings are based on nine publicly available data sets for seven different species. While the majority of these have sample sizes in the range 36–114, for completeness we also included two data sets consisting of 10 samples each. As the latter sample size does not lead to a powerful test of lognormality, it would be desirable to repeat the analysis with more extensive data once available. In addition, data from multiple subjects of the same species would allow testing the consistency of the lognormal distribution across individuals.

Finally, we propose a minimal model that accounts for the emerging lognormal distributions based on a noisy cell division process of the neural progenitor cells and their specification to neurons. In principle, a multiplicative process with Gaussian noise leads to lognormal distributions at any scale; the additive specification from progenitors to neurons approximately preserves the lognormality. However, this noisy process alone cannot explain the coexistence of lognormal distributions of *ρ* at different scales (within and across areas). When the distributed proliferation times are considered, the model can account for both within-area and across-area distributions.

The model explains the observed lognormality based on a minimal set of assumptions; hence, we did not include further mechanisms like cell death (apoptosis), migration, volumetric growth, the generation of other postmitotic cells, or area-specific proliferation rates. However, none of these additional mechanisms affect the main conclusions. Apoptosis is a widespread phenomenon (Inglis-Broadgate et al., 2005, Kalinichenko and Matveeva, 2008, Oppenheim, 1991) during neurogenesis and it can be modeled as a multiplicative process affecting the neuron density alone. Thus, the neuron density still depends linearly on the progenitor cell density and approximately maintains lognormality. While apoptosis could add to the inter-area variability, our model shows that it is not necessary for the distribution of variability per se—in agreement with Oppenheim et al. (1989) who found that the final pattern of spatial variability in spinal motoneuron density was already present before the onset of cell death and migration. Following the radial unit hypothesis (Rakic, 2009) we focus on radial migration of progenitor cells, ignoring tangential migration. In our model this assumption can be relaxed as long as there is no net increase or decrease in progenitor cell density due to tangential migration. Furthermore, we modeled the cell density in a fixed final target volume; thus, the effects of volumetric growth do not affect the model. The generation of other postmitotic cell types would reduce the effective proliferation rate of progenitor cells; this only leads to quantitative but not qualitative changes in the resulting distributions. Finally, it has been shown that cortical areas can have different proliferation rates (Dehay and Kennedy, 2007, Lukaszewicz et al., 2005, Polleux et al., 1997). If the proliferation rates are constant in time and lognormally distributed across areas, this additional variability would broaden the neuron density distribution but preserve the lognormal shape. However, in contrast to distributed proliferation times, it would not lead to the correlation between mean density and variability seen in the data (Figure 4C), because the proliferation rate affects the mean density but not the variance of the logarithmic progenitor density (see Eq. (7)). Furthermore, to the best of our knowledge, a difference in proliferation rate has only been shown for areas V1 and V2 (Dehay and Kennedy, 2007, Lukaszewicz et al., 2005)—thus, we speculate that this is an important reason for the drastically higher neuron density in V1.

In principle, cortex-wide organizational structures might be by-products of development or evolution that serve no computational function (Otopalik et al., 2017)— but the fact that we observe the same organizational principle for several species and across most cortical areas suggests that the lognormal distribution serves some purpose. Heterogeneous neuron densities could assist computation through their association with heterogeneity in other properties such as connectivity and neuronal time constants (Hilgetag et al., 2019, Rall, 1969); indeed, such heterogeneity is known to be a valuable asset for neural computation (Duarte and Morrison, 2019, Perez-Nieves et al., 2021). Alternatively, localized concentration of neurons in certain areas and regions could also serve a metabolic purpose since centralization might support efficient transport of metabolites across neurons and astrocytes (Bélanger et al., 2011, Magistretti and Allaman, 2015). Energy efficiency is particularly relevant since a large portion of the brain’s energy consumption is used to support the communication between neurons (Attwell and Laughlin, 2001, Laughlin and Sejnowski, 2003). Also from the perspective of cortical hierarchies it makes sense to have few areas with high neuron densities and many areas with lower neuron densities: Low-density areas contain neurons with large dendritic trees (Elston and Rosa, 1998) receiving convergent inputs from many neurons in high-density areas lower in the hierarchy. The neurons with extensive dendritic trees in higher areas are involved in different, area-specific abstractions of the low-level sensory information (Brincat et al., 2018, Kumar et al., 2007). All in all, there is probably not a single factor that leads to lognormal neuron densities in the cortex; further research will be needed to refine our findings and uncover the functional implications.

## Acknowledgements

This work was supported by the European Union Horizon 2020 Framework Programme for Research and Innovation (Human Brain Project SGA2 grant number 785907 and HBP SGA3 grant number 945539) and the Deutsche Forschungsgemeinschaft (DFG, German Research Foundation) under Priority Program (SPP 2041 “Computational Connectomics”) [S.J. van Albada: AL 2041/1-1 and 2041/1-2] and Open Access Publication Costs grant 491111487.

We thank Günther Palm for helpful discussions, Alexandre René for helpful discussions and support with the Bayesian model comparison, Anno Kurth for discussions about geometric Brownian motion, and Jon Martinez Corral for proofreading an early draft.

## Author contributions

Conceptualization AMG, AvM, SvA; Data curation AMG; Formal Analysis AMG, AvM; Funding acquisition SvA; Investigation AMG, AvM, SvA; Methodology AMG, AvM, SvA; Project administration SvA; Resources SvA; Software AMG, AvM; Supervision SvA; Validation AvM; Visualization AMG; Writing original draft AMG, AvM; Writing – review & editing AMG, AvM, SvA.

## Competing interests

The authors declare no competing interests.

## Data and materials availability

This work produced no new data and instead relied on a corpus of neuron density data available from the literature, which we gratefully acknowledge; see Cell density data for a detailed description.

## Supplementary tables

**Table S1:**
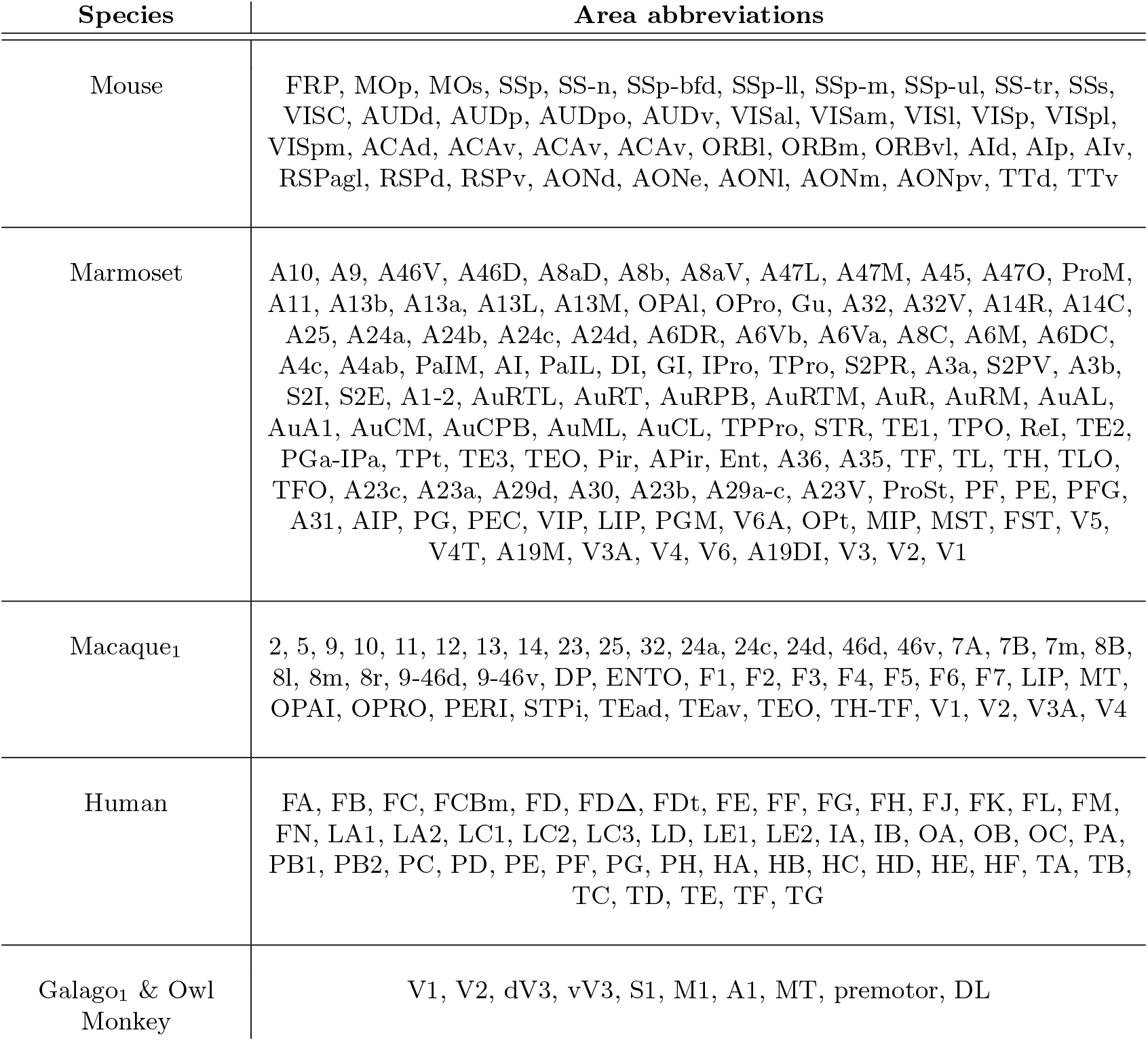
Cortical areas included in this study.

**Table S2:**
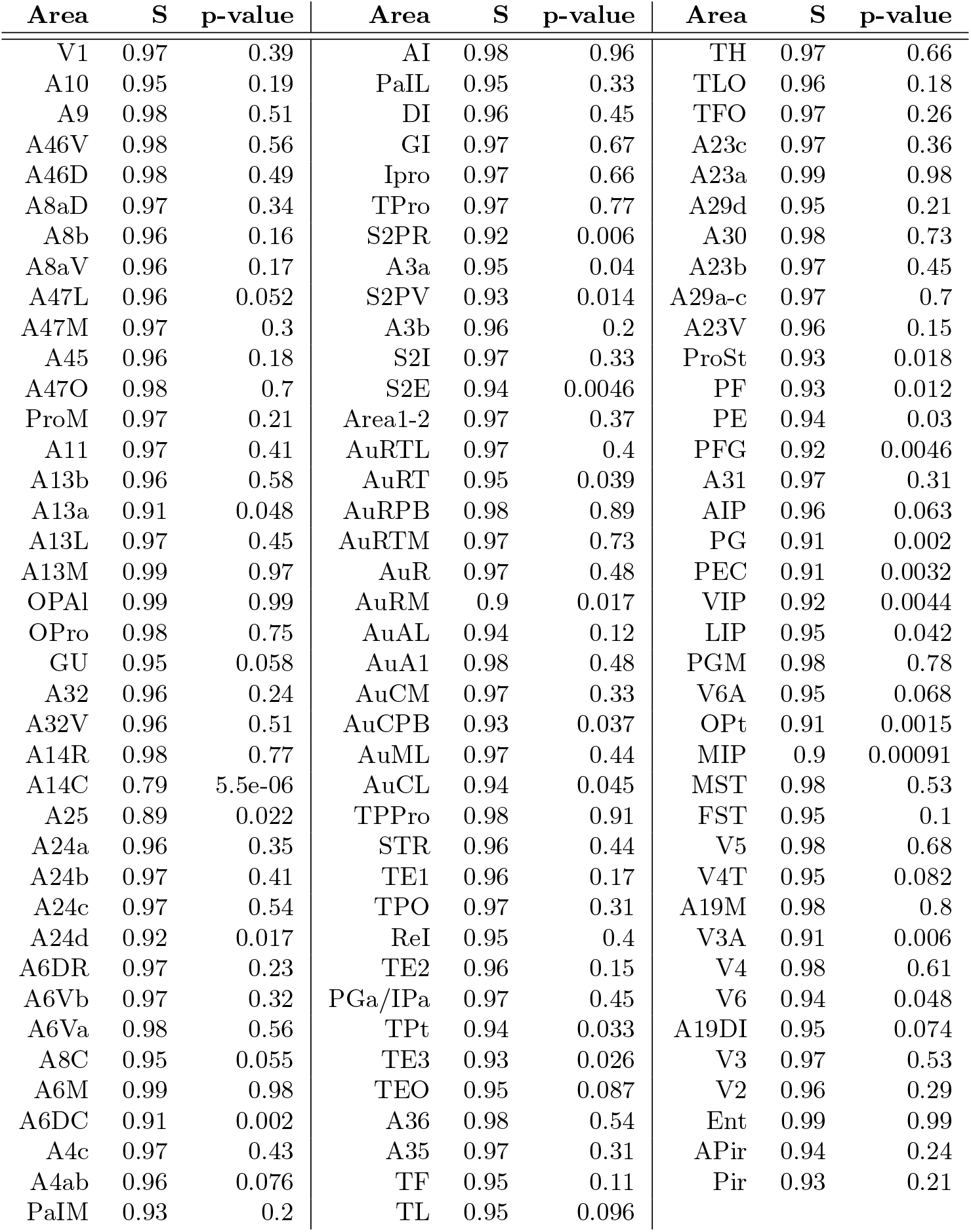
Results of the Shapiro-Wilk test for normality of ln(*ρ*_*s*_) in marmoset cortical areas. Values rounded to two significant digits.

## Supplementary figures

**Figure S1:**
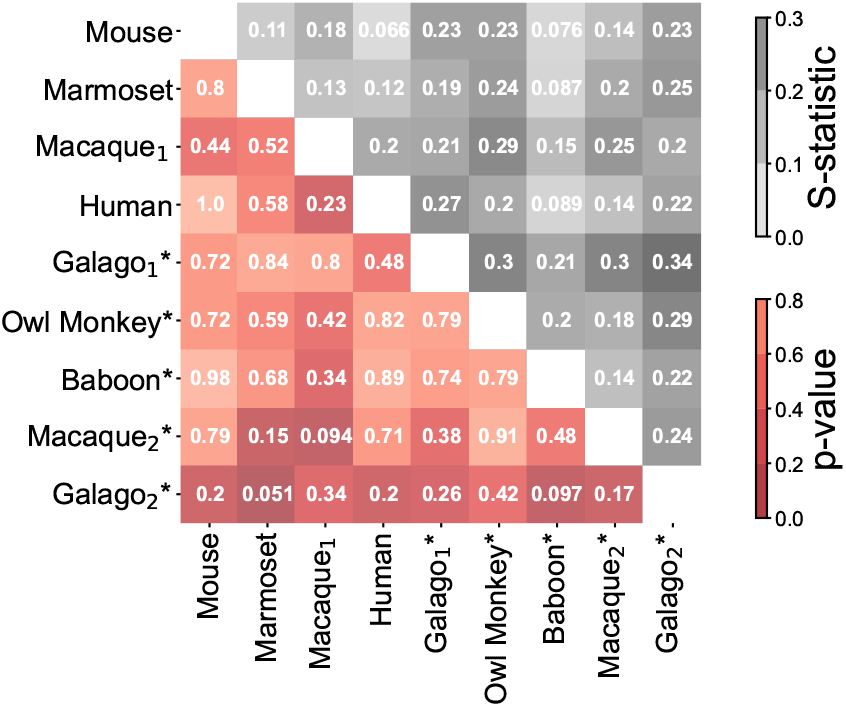
Pairwise Kolmogorov-Smirnov two-sample two-sided tests. P-values and S-statistics displayed below and above the diagonal, respectively. The z-scored log neuron density distributions of the four species are statistically indistinguishable at the 0.05 level.

**Figure S2:**
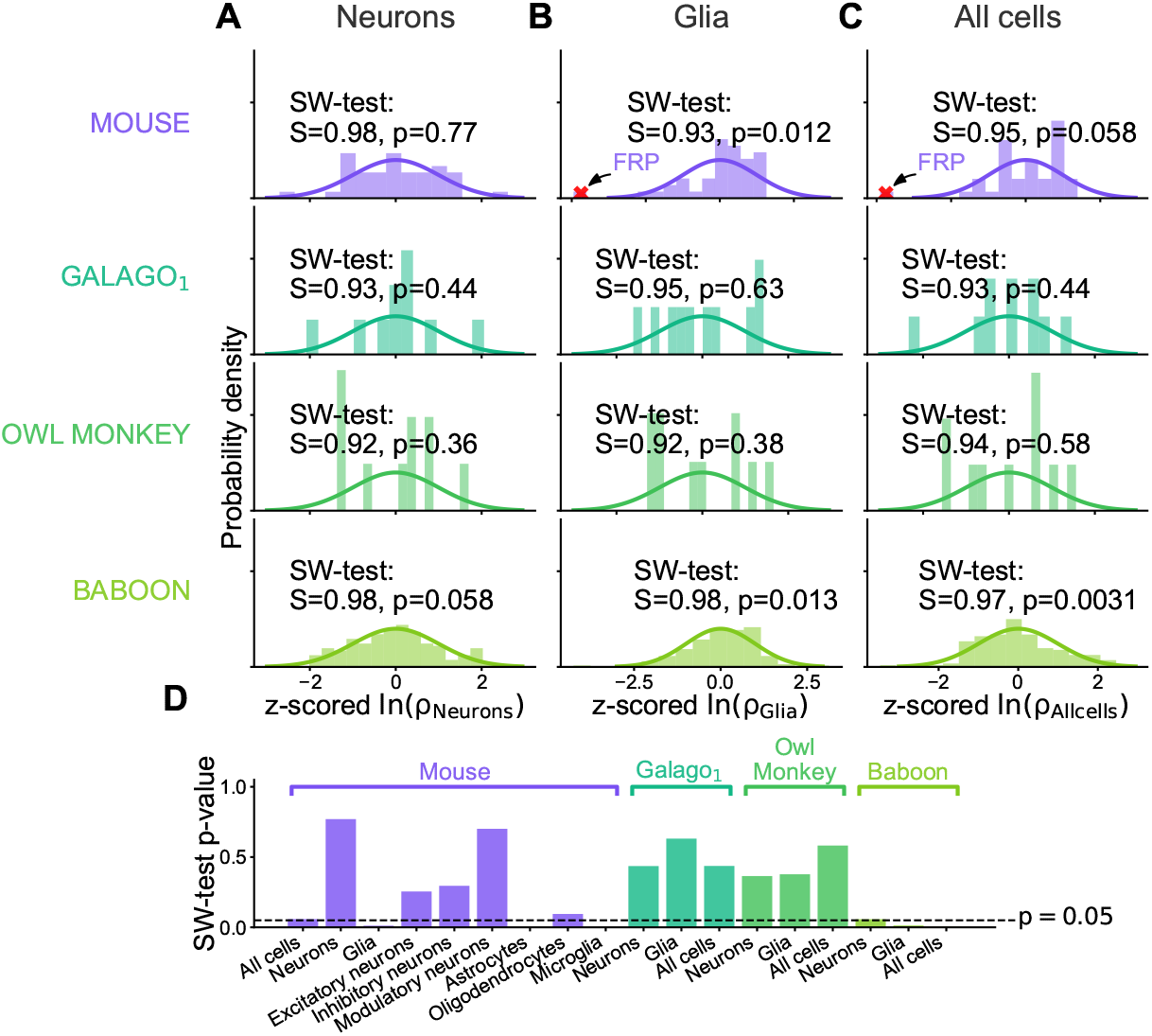
Comparison of neuron and glia lognormality. **A**–**C** Histogram of z-scored log density and result of Shapiro-Wilk test for neurons (**A**), glia (**B**), and all cells combined (**C**). **D** Barplot of p-values resulting from Shapiro-Wilk normality test for all cell types. Panel **A** is equivalent to the data shown in Figure 1.

**Figure S3:**
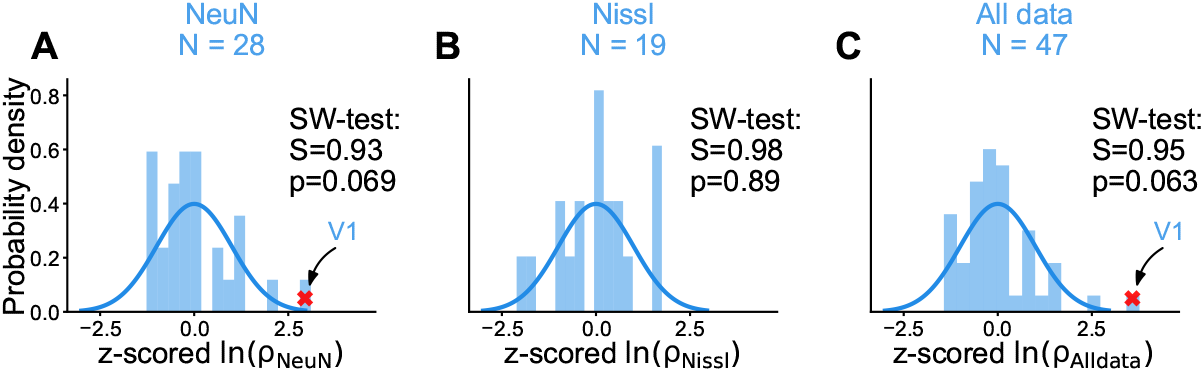
Lognormality of cell densities from different staining types in macaque cortex based on the macaque_1_ data set. **A**-**C** Histogram of z-scored log density and result of Shapiro-Wilk test for NeuN staining only (**A**), Nissl staining only (**B**) and all measurements combined (**C**). The Nissl data were scaled down based on the linear relationship with the NeuN data (Beul and Hilgetag, 2019). Red crosses indicate outliers (| z-scored ln(*ρ*)| ≥ 3, which were excluded from the test. Panel **C** is equivalent to the data shown in Figure 1.

**Figure S4:**
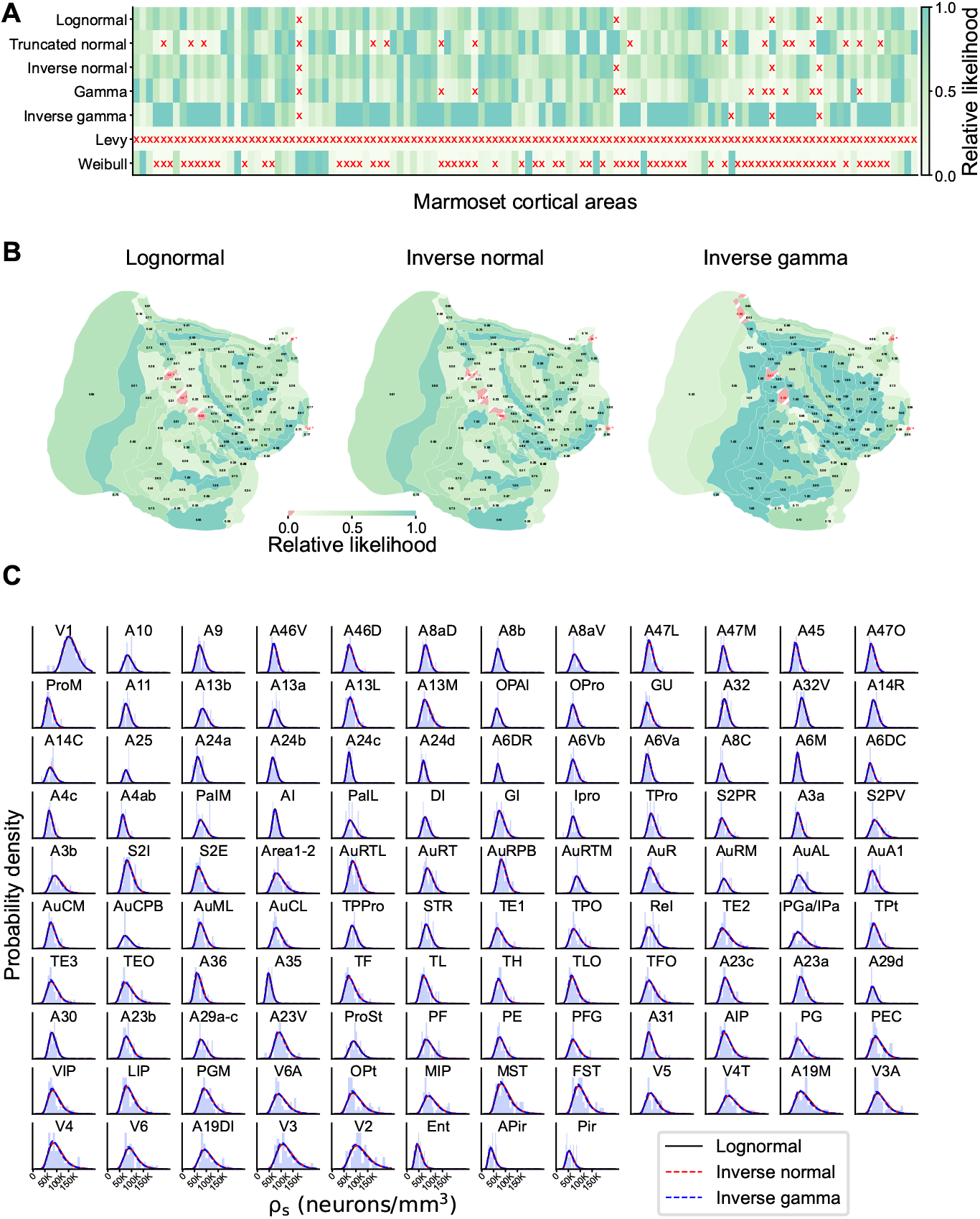
Statistical model comparison within the marmoset cortical areas. **A** Relative likelihood for seven compatible statistical models for all areas of the marmoset; a red cross (x) indicates a relative likelihood < 0.05 with respect to the model with the highest likelihood. **B** Spatial distribution of relative likelihood for the three best statistical models. **C** The three best statistical models fitted to the neuron density histograms in each area of marmoset cortex; the three models produce visually nearly indistinguishable fits.

**Figure S5:**
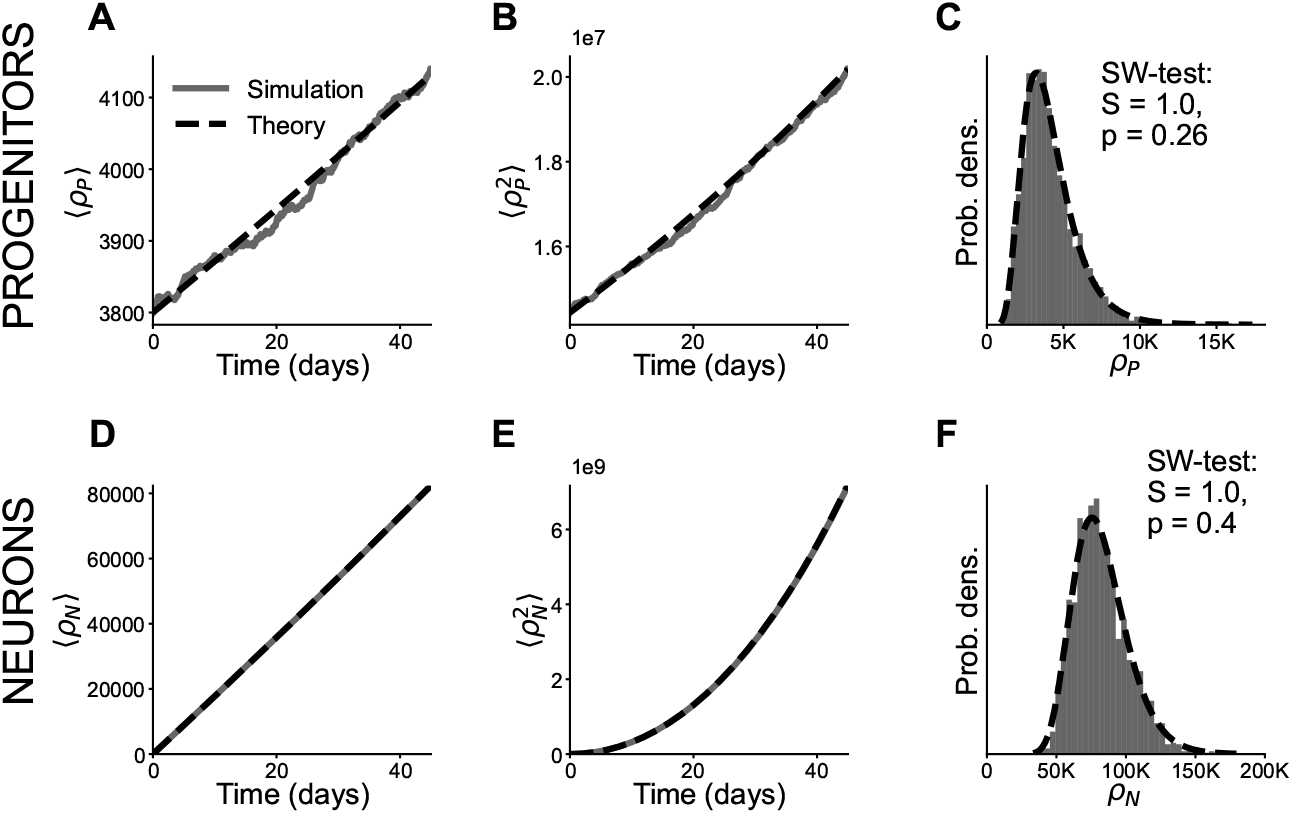
Verification of neurogenesis model. **A, B, D, E** Time evolution of the mean density of progenitors (A) and neurons (D) and the corresponding second moment for progenitors (B) and neurons (E). Theory based on Equations (8) and (10). **C, F** Resulting progenitor (C) and neuron (F) density distributions at the end of neurogenesis. Both distributions are compatible with a lognormal; for the neuron density this is not a formal result but rather an approximation based on Equation (11). Simulation parameters: time step Δ*t* = 0.05 days, total time *T* = 45 days, *N*_sample_ = 2000 samples, initial progenitor density *ρ*_0_ = 3.8 × 10^3^, average rate *λ* = 2 log(2)*/*3 days, and noise intensity *σ* = 0.061 days^−1*/*2^.

## Notes

### Competing Interest Statement

The authors have declared no competing interest.

### Summary of Updates

Reordered content, extended details in methods section. Moved some results away from the methods into the results section.

